# Online spike sorting via deep contractive autoencoder

**DOI:** 10.1101/2021.04.23.441225

**Authors:** Mohammadreza Radmanesh, Ahmad Asgharian Rezaei, Alireza Hashemi, Mahdi Jalili, Morteza Moazami Goudarzi

## Abstract

Spike sorting – the process of separating spikes from different neurons – is often the first and most critical step in the neural data analysis pipeline. Spike-sorting techniques isolate a single neuron’s activity from background electrical noise based on the shapes of the waveforms (WFs) obtained from extracellular recordings. Despite several advancements in this area, an important remaining challenge in neuroscience is online spike sorting, which has the potential to significantly advance basic neuroscience research and the clinical setting by providing the means to produce real-time perturbations of neurons via closed-loop control. Current approaches to online spike sorting are not fully automated, are computationally expensive and are often outperformed by offline approaches. In this paper, we present a novel algorithm for fast and robust online classification of single neuron activity. This algorithm is based on a deep contractive autoencoder (DCAE) architecture. DCAEs are deep neural networks that can learn a latent state representation of their inputs. The main advantage of DCAE approaches is that they are less sensitive to noise (i.e., small perturbations in their inputs). We therefore reasoned that they can form the basis for robust online spike sorting algorithms. Overall, our DCAE-based online spike sorting algorithm achieves over 90% accuracy at sorting previously-unseen spike waveforms. Moreover, our approach produces superior results compared to several state-of-the-art offline spike-sorting procedures.

## Introduction

Spike sorting refers to the process of isolating spikes according to the individual neurons that generate them. It presents a significant challenge in neural data analysis [1], and consists of three major steps: filtering, detection, and clustering, each of which critically affects spike sorting accuracy. Conventional approaches to spike sorting consist of four components: 1) Band-pass filtering of raw signals, 2) Extracting relevant waveform features, 3) Feeding these features to a clustering algorithm, and 4) Using these clusters as inputs to a classifier. The classifier associates each cluster with the activity of a single neuron. The approach we outline in this paper consists of these four components (Fig. 1).

**Figure 1:**
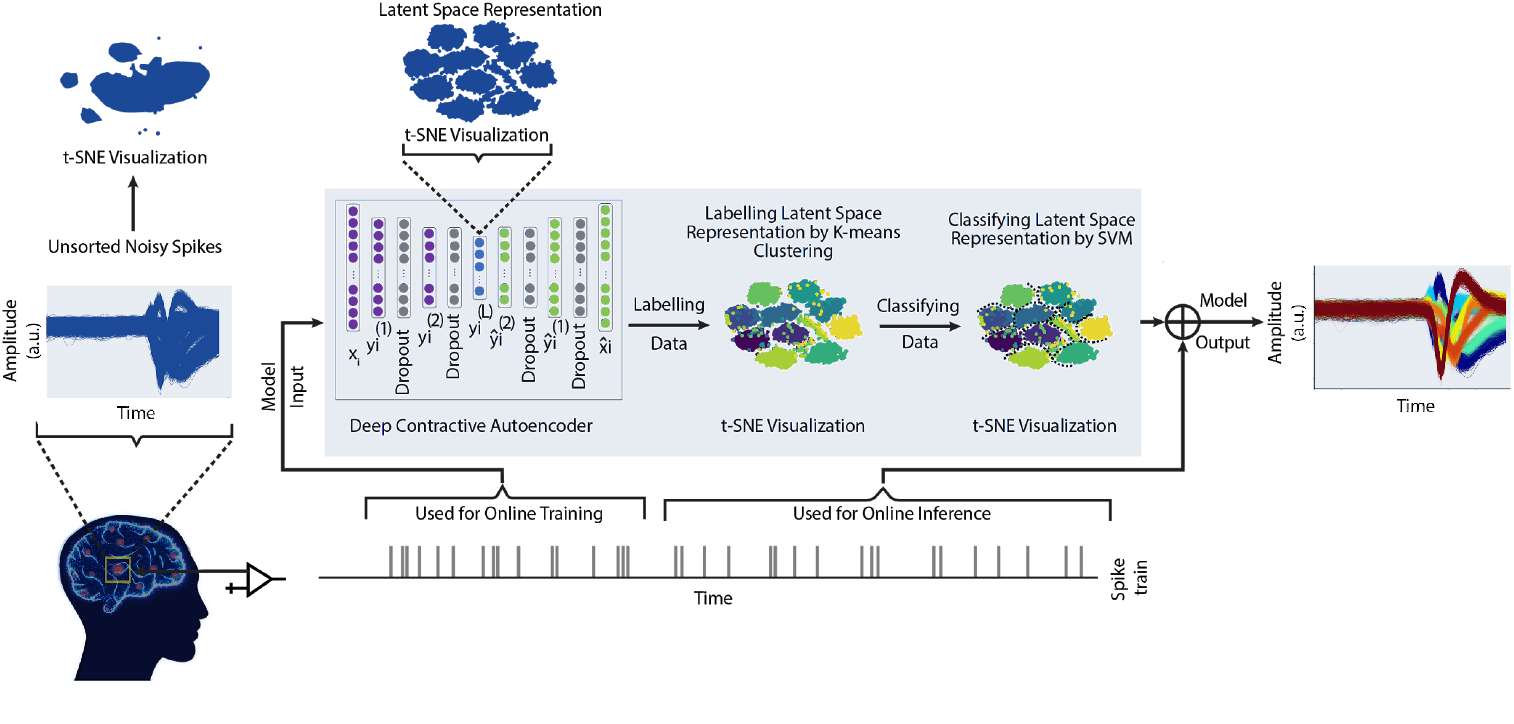
Different stages of the spike-sorting algorithm. Our end-to-end method consists of three main components: First, a contractive autoencoder which consists of two hidden layers for both the encoder and decoder layers of our neural network. A dropout layer immediately follows each hidden layer in order to prevent overfitting. The output of the latent space representation produced by the autoencoder (blue nodes) is fed into a K-means clustering algorithm to label the embedded samples in an unsupervised manner.

Efficient and reliable spike sorting is an ever more crucial problem for neuroscience practitioners. The number of neurons that can be recorded simultaneously has doubled every 7 years [2], while advancements in spike sorting have lagged considerably. This gap between the capacity of neural recording devices and the limits of current spike-sorting methods is growing even larger. Existing algorithms are simply unable to cope with the challenge of processing such high volumes of data.

To address this gap, more recent algorithms have significantly improved spike sorting procedures, greatly reducing the need for manual intervention [3]. These methods have focused on improving spike detection through the use of template-based filtering [4, 5], wavelet transforms [6] and energy operators [7]. However, most of these approaches are ‘offline’, i.e. their application is limited to cases where spikes are sorted after the data has already been acquired. In contrast, the novel algorithm that we introduce here can be adapted to online (as well as offline) applications. Apart from introducing a fast and effective means of sorting spikes on the fly, we also propose an algorithm that is robust to both measurement noise, and noise due to electrode drift (a significant challenge for spike sorting using chronically implanted electrodes).

While different components of every step in the spike sorting pipeline are the subject of active research, our approach focuses on robust feature extraction. Inspired by the design of denoising [8] and contractive auto-encoders [9], we propose an embedding method for learning a noiseless low dimensional representation from input waveforms. Although the idea of learning a low dimensional representation from extracellular recordings has been implemented in previous work [8], these algorithms often do not take the presence of background noise in the raw spikes into consideration. In contrast, the tests we ran on our models unequivocally demonstrate that our proposed method is robust to different levels of background noise, while the performance of other state-of-the-art models drops as we introduce additional background noise into the raw signal.

## Results

As mentioned, our proposed end-to-end model consists of three different components or processes: a deep contractive autoencoder (DCAE), which is followed by clustering and classification procedures (Fig. 1). The DCAE takes in high dimensional data as its input and maps it onto a low-dimensional latent space that generates the inputs for subsequent stages of the spike-sorting pipeline. We visually demonstrate the robustness of this approach by injecting different levels of additive noise into a sample spike waveform (Fig. 2). We demonstrate that DCAE can produce near-identical low-level representations of the waveform for different amounts of injected noise.

**Figure 2.**
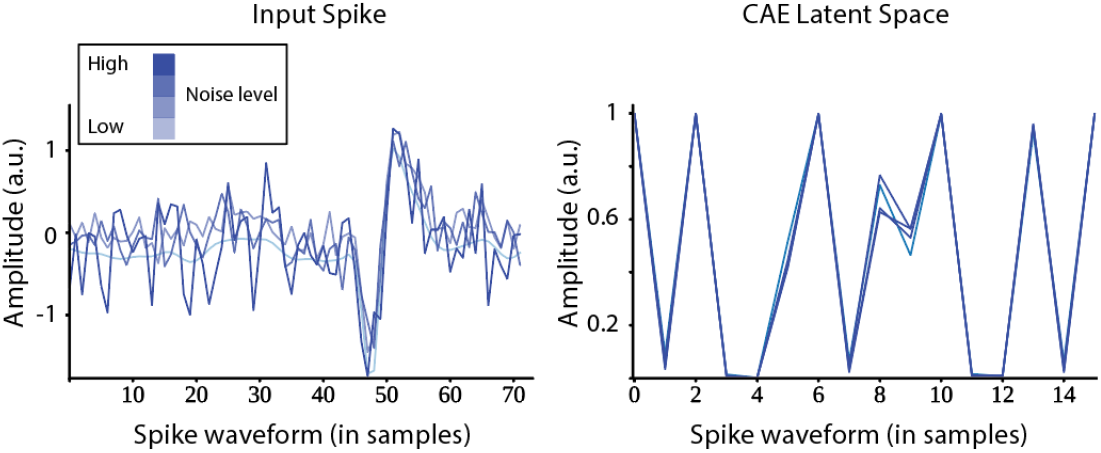
DCAE with additive noise. (a) Example waveform. We increased the amount of additive noise injected into this waveform (light to dark blue). (b) Latent space representation of the example waveform. The latent space representation was near identical for all noise conditions (light to dark blue).

Furthermore, we compare the performance of our algorithm against several others when different sizes of training sets are used. We also evaluate our model’s robustness relative to a standard autoencoder by applying different levels of noise to our training set. Finally, we use gap statistics to demonstrate that there is no need to pre-specify the exact number of clusters for our proposed algorithm. Our comparisons were made across four simulated ground-truth extracellular recordings (described below).

We used four publicly available datasets in this study [10].

- Neurocube_Easy1, Neurocube_Easy2, and Neurocube_Sim2: These datasets have been adapted from the SYNTH_MONOTRODE dataset [10]. All of them generate spike waveform templates based on single-neuron simulations, and randomly placing the spike waveforms conforming to pre-specified ISI distributions. These datasets consist of a single channel, but with a different number of units Neurocube_Easy1 and Neurocube_Easy2 have an intrinsic noise level of 0.005, whereas Neurocube_Sim2 is noise free.
- SpikeInterface_Synthetic: This is a single channel dataset consisting of 10 units generated using the SpikeForest platform.

Additional details about the datasets we used are presented in the following table (Table 1).

**Table 1:** Statistics of Synthetic Electrophysiology Recordings.

|  | Sample Rate (KHz) | # Channels | Duration (sec) | # Units |
| --- | --- | --- | --- | --- |
| Neurocube_Easy1 | 24 | 1 | 60 | 3 |
| Neurocube_Easy2 | 24 | 1 | 60 | 3 |
| Neurocube_Sim2 | 24 | 1 | 600 | 16 |
| SpikeInterface_Synthetic | 24 | 1 | 4000 | 10 |

As demonstrated in Fig. 2, when dealing with intrinsically noisy data, adding a contractive loss term to a standard autoencoder leads to the generation of more robust representations. We specifically consider two alternative forms of contractive autoencoders in our study: 1-Semi-supervised and 2-Unsupervised (see Methods).

The main difference in these approaches lies in the fact that the semi-supervised online CAE (hereafter referred to as SOCAE) relies on ground truth labels, whereas unsupervised online CAE (UOCAE) uses labels derived from a K-means algorithm. As such, UOCAE produces slightly lower-accuracy values relative to SOCAE. Nonetheless, UOCAE substantially outperforms (both in terms of accuracy and runtime) other SOTA spike sorters, and a support vector machine (SVM) classifier that was trained on the original dataset (Fig. 3). Importantly, the accuracy values for both UOCAE and SOCAE remain largely unaffected by training dataset size, implying that DCAE is an appropriate algorithm for online spike sorting, given that it can produce reliable results on small datasets.

**Figure 3:**
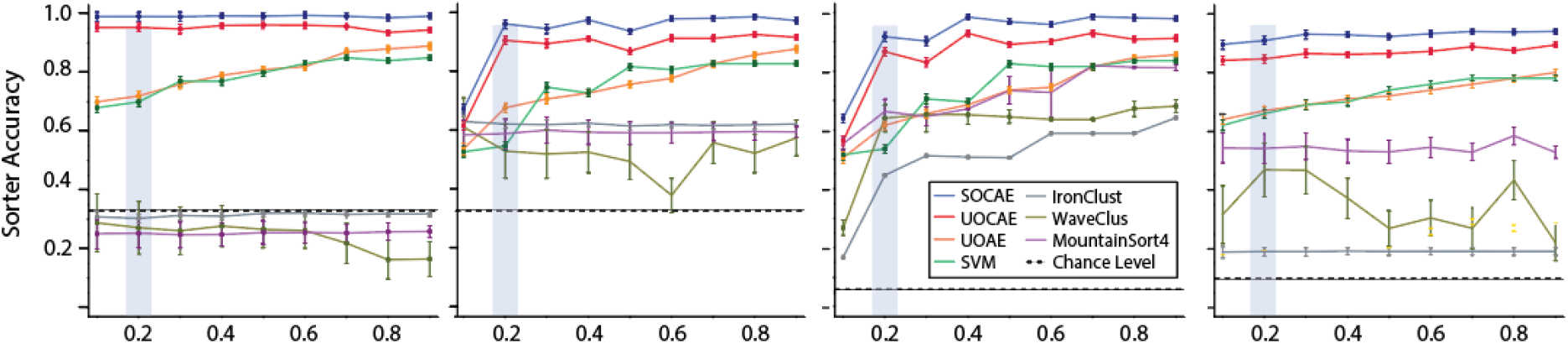
Spike sorting accuracy as a function of training size. Our proposed algorithm produced much higher accuracy scores than the other alternative we tested.

We further assessed the superiority of UOCAE over alternative spike sorting methods using several standard performance metrics, such as accuracy, precision, and recall (Table 2). Taken together, these measures fully characterize the efficacy of any classification algorithm. We demonstrate that, out of all the standard spike-sorting algorithms we evaluated, MountainSort4 achieved the highest accuracy, nonetheless our DCAE algorithm vastly outperformed this algorithm.

**Table 2:** Summary of accuracy metrics for the algorithms we tested on different datasets.

|  | Methods | DCAE Sorter |  |  | MountainSort4 |  |  | IronClust |  |  | WaveClus |  |  |
| --- | --- | --- | --- | --- | --- | --- | --- | --- | --- | --- | --- | --- | --- |
|  |  | Accuracy | Recall | Precision | Accuracy | Recall | Precision | Accuracy | Recall | Precision | Accuracy | Recall | Precision |
| Neurocube_Easy1 |  | 0.97 | 0.99 | 0.98 | 0.26 | 0.28 | 0.3 | 0.31 | 0.32 | 0.33 | 0.21 | 0.2 | 0.26 |
| Neurocube_Easy2 |  | 0.96 | 0.99 | 0.96 | 0.6 | 0.62 | 0.63 | 0.62 | 0.64 | 0.65 | 0.51 | 0.51 | 0.6 |
| Neurocube_Sim2 |  | 0.91 | 0.98 | 0.93 | 0.81 | 0.84 | 0.84 | 0.64 | 0.67 | 0.66 | 0.69 | 0.73 | 0.7 |
| Neurocube_Synt |  | 0.87 | 0.94 | 0.92 | 0.58 | 0.64 | 0.7 | 0.19 | 0.2 | 0.19 | 0.35 | 0.37 | 0.4 |

We further sought to demonstrate the extent to which our model was robust to additive noise. We applied different levels of noise to our training dataset and evaluated the effects of injecting increasingly large quantities of noise into our training datasets on classifier accuracy. We found that DCAES (compromising both UOCAE and SOCAE) achieved similar rates of classifier accuracy in spite of the different levels of noise injected into the training dataset. This could be contrasted with UOAE whose performance dropped as a function of increasing noise levels. These results serve to further demonstrate the robustness of DCAE models in spike sorting applications, relative to standard autoencoders.

A major challenge in cluster analysis is determining the optimal number of clusters in a particular dataset. The gap statistic is a standard unsupervised method for overcoming this challenge. As such, we sought to establish that our DCAE methods can automatically converge on the correct (“ground-truth”) number of clusters by adopting this method. As Fig. 5 illustrates, our model could identify the exact number of clusters consistently across all datasets. In other words, we found that our algorithm did not need to have access to the correct number of clusters in advance in order to identify them accurately.

**Figure 4:**
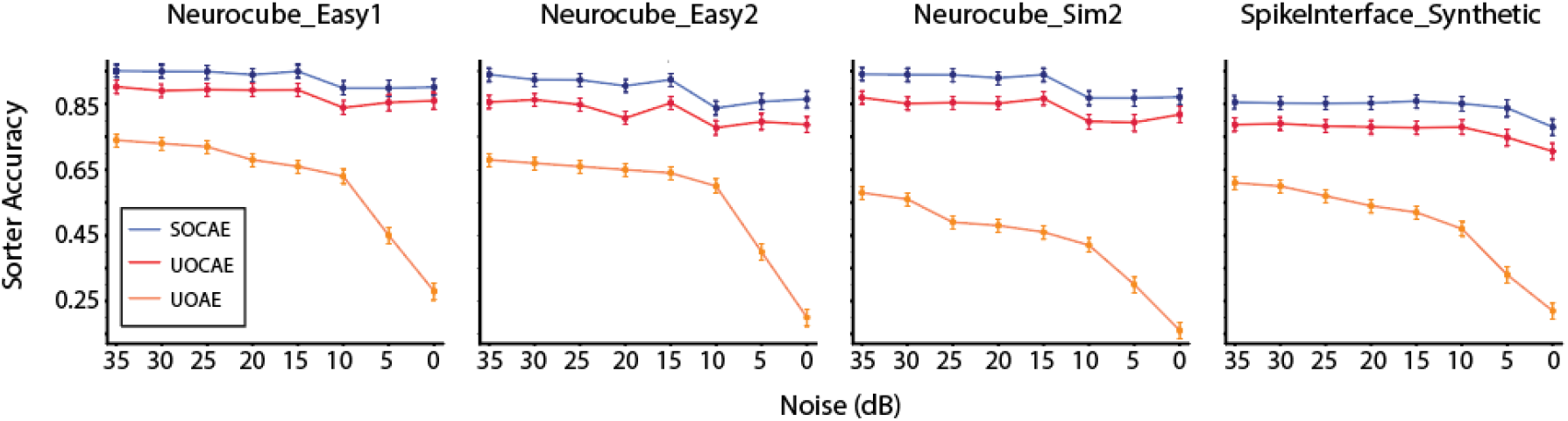
Sorter accuracy results on different levels of noise. To show our model’s robustness to noise, different levels of noise were applied to 20% of the training data.

**Figure 5:**
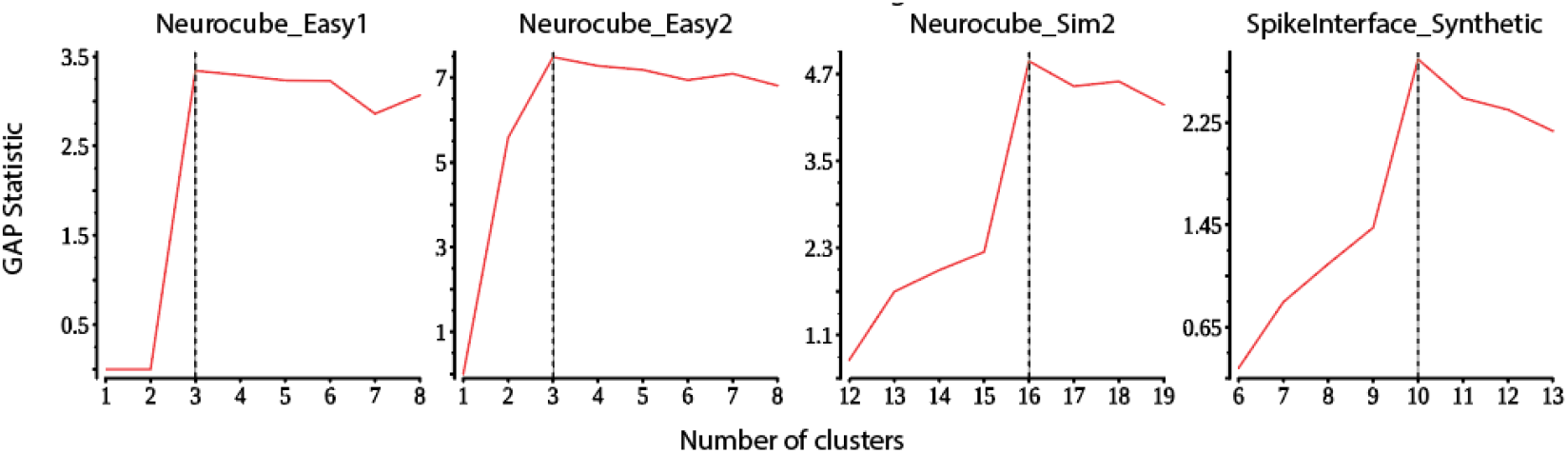
Gap statistic results. Our model with a Gap statistic determines the correct number of clusters automatically.

**Figure 6:**
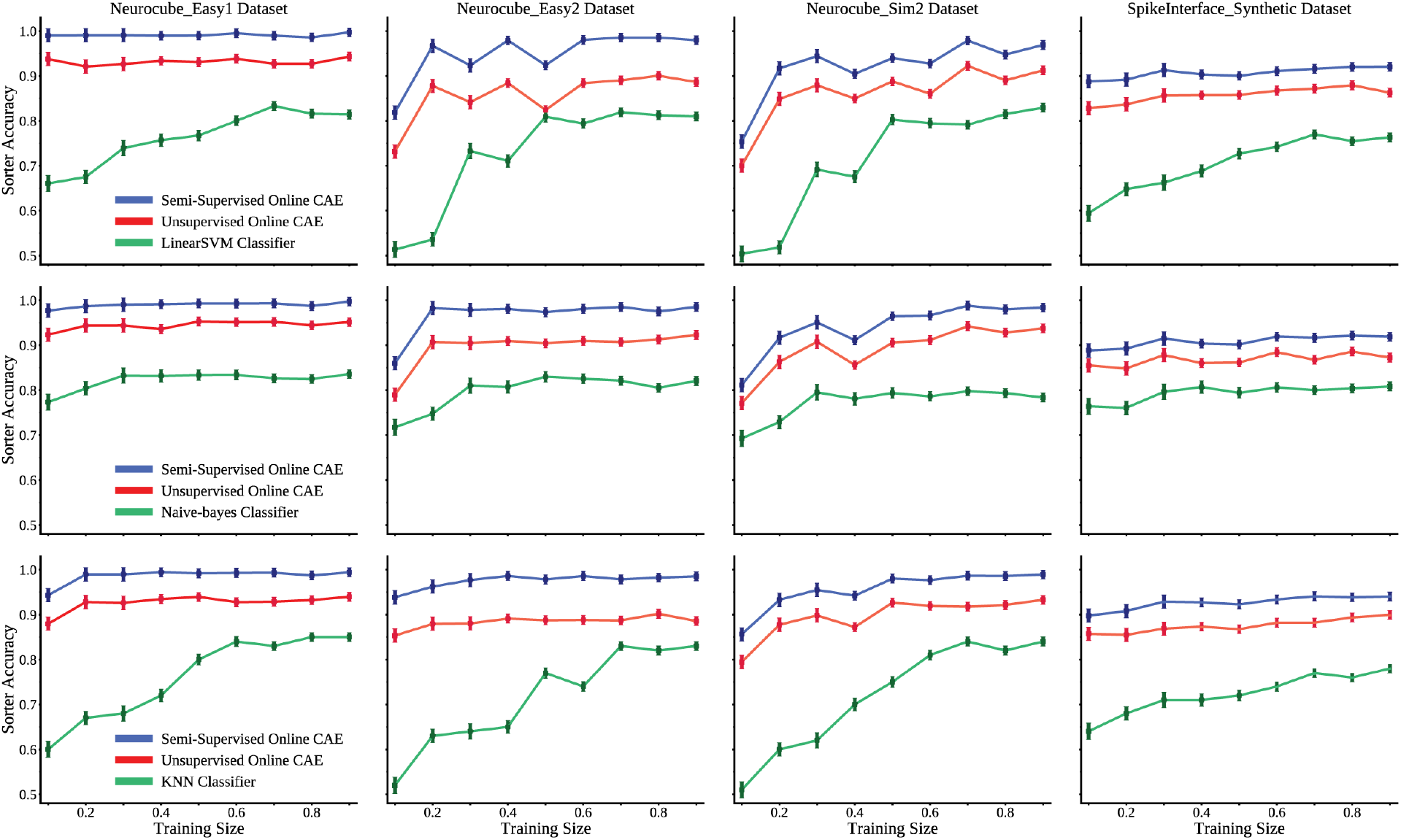
Spike sorting accuracy for different classifiers as a function of training dataset size. Our DCAE algorithms outperformed three different vanilla classifiers (Linear SVM, Naive Bayes and k-NN).

Finally, we compared and contrasted the performance of our DCAE algorithms by substituting the SVM classifier with three other basic classifiers: 1) linear SVM, 2) Naive Bayes and 3) K-nearest neighbours. We refer to these classifiers as “basic” because they are among the most common out of the box procedures used in supervised learning. We demonstrate that neither of these basic classifiers is better at sorting spikes than an algorithm that combines the same basic classifier with a DCAE (be it SOCAE or UOCAE), and while the performance of the basic classifiers improved by increasing the size of the training dataset, it was never as high as any of our DCAE algorithms combined with a basic classifier.

## Discussion

In this paper we introduced a novel algorithm for spike sorting and validated it using four synthetic datasets. We demonstrated three key properties for our DCAE algorithms: 1) They produce representations of the data that are robust to additive noise. We demonstrated that the performance of our algorithm did not significantly drop with the addition of noise, in contrast to other traditional methods. 2) They outperform several SOTA models in various online and offline spike-sorting applications. 3) They reliably classify spikes for small and large datasets.

One shortcoming of this approach is that we restricted our evaluation of DCAE to datasets that consisted of a single channel. In the future we wish to apply the same algorithm to datasets consisting of multiple channels, as we believe that it will achieve similarly high levels of accuracy for such datsets.

## Method

A novel approach for regularizing auto-encoders has recently been proposed, termed Deep Contractive Auto-Encoders (DCAE) [9]. It shares a similar motivation to DAEs in that it aims to be robust to small variations in the training dataset [8]. However, DCAE achieves its robustness in a rather different manner: instead of stochastically corrupting the input, it balances the reconstruction error with an analytical penalty term that penalizes the Frobenius norm of the encoder’s Jacobian at training points. In doing so, DCAE algorithms seek to generate representations that are robust to noise, whereas DAEs are concerned with generating a noiseless reconstruction of their inputs.

The basic Auto-Encoder (AE) framework considered here starts from an encoding function *f* that maps an input 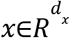 to a hidden representation 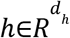 where 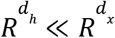. It has the form:

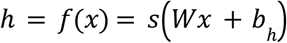

where *s* is the logistic sigmoid activation function 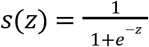. The encoder is parametrized by *d_h_* × *d_x_* weight matrix *W*, and a bias vector 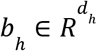.

A decoder function *g* then maps the hidden representation *h* back to a reconstruction *y* as follows:

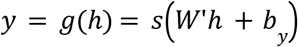

The decoder’s parameters are a bias vector 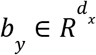, and a matrix *W*’. In this paper we only explore the case in which *W*’ = *W*^*T*^.

Basic auto-encoder training consists of finding parameters θ = {*W, b*_*h*_, *b*_*y*_} that minimize the reconstruction error on a training set of examples *D*_*n*_, i.e. minimizing the following objective function:

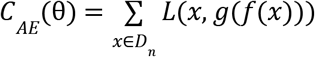

where *L*(*t, r*) is the reconstruction error between target *t* and reconstruction *r* (typically a squared error or cross-entropy loss).

To encourage *f*(*x*) to be robust to small variations of training input *x*, we penalize its sensitivity to that input, measured as the Frobenius norm of the Jacobian, 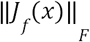.

A CAE is trained to optimize the following objective function:

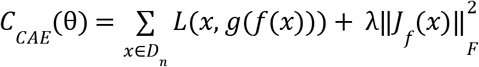

where λ is a positive hyperparameter that controls the strength of the regularization. One can see that:

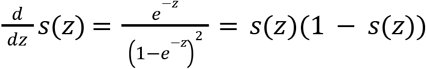

Where *h* = *f*(*x*), the linear + sigmoid mapping yields a simple expression for the penalty term [9]:

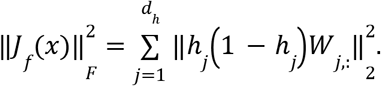

## Author contributions

M.M.G: Conceptualization, Methodology, Software, Visualization, Writing, Supervision. M.R.: Methodology, Formal Analysis, Software, Visualization, Writing. A.A.R.: Software, Writing. A.H.: Visualization, Writing. M.J.: Writing., Supervision.

